# Integrating Multimodal Data for a Comprehensive Knowledge Graph to Advance Infectious Disease Research

**DOI:** 10.1101/2025.06.02.657361

**Authors:** Hengyu Fan, Liwei Guo, Fang Li, Zhen Yuan, Yuxi Deng, Yingjing Xiao, Honglin Li, Shiliang Li

**Affiliations:** Innovation Center for AI and Drug Discovery, School of Pharmacy, East China Normal University, Shanghai 200062, China; School of Data Science and Engineering, East China Normal University, Shanghai 200062, China; Shanghai Key Laboratory of New Drug Design, School of Pharmacy, East China University of Science and Technology, Shanghai 200237, China; School of Life Sciences, East China Normal University, Shanghai 200241, China; School of Computer Science and Technology, East China Normal University, Shanghai 200062, China; Department of Pain management, HuaDong Hospital affiliated to Fudan University, Shanghai, 200040, China

**Keywords:** Knowledge graph, Infectious disease, Drug repurposing, Natural language processing

## Abstract

Infectious diseases remain a formidable threat to global public health, with their escalating morbidity and mortality rates compounded by recurrent epidemics and the alarming rise of antimicrobial resistance (AMR). These challenges have intensified the urgent demand for innovative therapeutic strategies that can accelerate drug development cycles and overcome traditional research bottlenecks. To address these critical needs, we present IDKG (Infectious Disease Knowledge Graph), a specialized large-scale biomedical knowledge network designed to bridge data fragmentation through multimodal data integration. The IDKG constructs comprehensive associations from 345 infectious diseases and 708 pathogens across heterogeneous biomedical sources systematically. The graph architecture comprises nearly 50,000 nodes (8 types, including Pathogen, Protein, etc.) and over 1.2 million edges (11 types, including treats, contains, etc.), establishing an interconnected framework that enables systematic interrogation of cross-disciplinary knowledge. The integrative approach effectively dismantles conventional data silos while preserving biological contextuality. We validated the IDKG’s potential by applying graph neural network-based approaches for drug repurposing prediction in human metapneumovirus (hMPV) infection, a common acute respiratory infection for which effective specific antiviral drugs are currently absent. The successfully identification of established antiviral agents, such as ribavirin and emetine, by our M1 model demonstrated its predictive accuracy and biological relevance. IDKG unifies multimodal biomedical data into a network to accelerate drug discovery and bolster outbreak response. This establishes a data-driven, knowledge-based paradigm for infectious disease research.

**Graphical Abstract:** 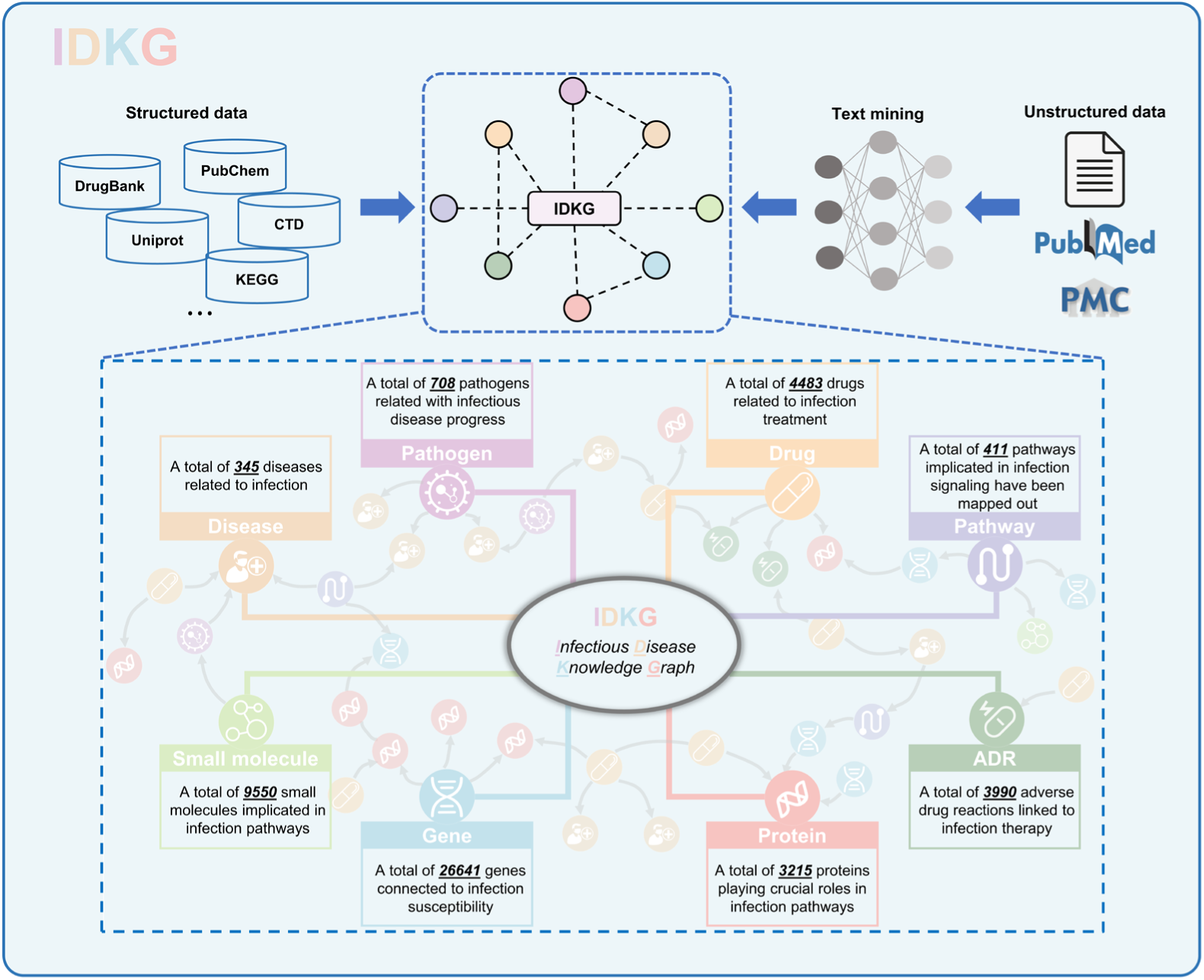

## Introduction

Infectious diseases remain significant global health threats due to their complexity and rapid spread, presenting formidable challenges to public health systems worldwide^1^. The devasting impact of pandemics has been repeatedly demonstrated throughout history^2^. Studies estimate that the COVID-19 pandemic resulted in a staggering 18.2 million deaths globally within its first two years^3^. Beyond pandemics, antimicrobial resistance (AMR) has emerged as one of the most pressing public health crises, urgently requiring innovative and effective solutions^4,5^. A report by the Global Leaders Group on Antimicrobial Resistance predicts that without targeted efforts to combat AMR, global life expectancy could decrease by 1.8 years within the next decade^6,7^. Despite this growing threat, the development pipeline for antibiotics remains critically insufficient, failing to meet both current and future demands^8^. Over the past two decades, the number of newly approved antibiotics has decreased sharply, even as multidrug-resistant pathogens continue to rise^9^. Exacerbating this challenge is the substantial financial costs and prolonged timelines required for developing clinically viable drugs^10,11^. These compounding issues highlight the urgent need for innovative and time-efficient strategies to accelerate drug discovery and address the critical demand for effective antimicrobial therapies.

In the biomedical domain, big data is revolutionizing traditional paradigms by integrating multisource information from laboratory experiments, clinical studies, medical records, and the Internet of Medical Things (IoMT). Omics technologies, such as genomics and proteomics, have dramatically enhanced high-throughput data acquisition efficiency^12^. However, the exponential growth of data volume has exacerbated complexities in biomedical research—heterogeneous data formats across domains and systemic interoperability barriers lead to persistent data silos, hindering comprehensive knowledge discovery and translational applications across multidimensional datasets^12,13^. In the yield of infectious diseases, characterized by their dynamic nature and inherent complexity^14^, the challenges become particularly pronounced. This necessitates the development of tools capable of integrating multimodal data to uncover latent associations and novel insights. Knowledge graph (KG) offers a structured and efficient way to address these challenges.

KG is a graph representation in which information entities are represented as nodes, and their relations are coded as edges connecting the corresponding nodes^15^. This structure enables the explicit integration of heterogeneous data sources while preserving the intricate relationships essential for biomedical research. Knowledge graphs have been proven^16,17^ to be an effective representation for knowledge inference. A wide adoption of KG-based deep learning (DL) methods has led to significant advances in the field of drug discovery, especially in drug repurposing and drug target discovery^18–21^. Based on the extensive knowledge and topological structure of KGs, DL methods can learn potential features from highly complex biomedical networks to improve predictive performance in relative research.

While general-purpose KGs have demonstrated utility in biomedical applications, including clinical decision support, drug discovery and treatment recommendation^22^, infectious diseases present unique and complex challenges that existing KGs fail to adequately address. The rapid evolution of pathogens, the rise of AMR, and the intricate host-pathogen interactions demand a domain-specific approach. Infectious diseases, unlike many other therapeutic areas, involve multifaceted biological processes that require the integration of cross-disciplinary information such as epidemiologic data and pathogen specific targets.

In response to these challenges, we constructed a multidimensional biomedical knowledge graph of infectious disease (IDKG). Our pipeline involved processing structured databases, extracting information from unstructured literature through text mining, and integrating multimodal data into a standardized graph-based framework. The IDKG is a massive resource, comprising nearly 50,000 nodes (8 types) and a vast network of over 1.2 million edges (11 types), encompassing 345 infectious diseases and 708 pathogens. By unifying heterogeneous data, the IDKG systematically unveils the intricate relationships among various entities, and holds potential for drug repurposing when coupled with GNN-based reasoning algorithms. This valuable resource is poised to empower researchers in advancing drug development and proposing innovative therapeutic strategies.

## Results

### Statistical data of IDKG

The IDKG was constructed by integrating a wide range of multimodal data sources, including infectious disease related databases and literature, biopharmaceutical resources like DrugBank and UniProt, and insights mined from unstructured text (Figure 1). The data gathered included a total of 49,343 nodes and 1,231,991 edges which curated 345 infectious diseases and 708 pathogens. This represents, to our knowledge, the largest knowledge graph focused on infectious diseases. We classified the nodes into 8 main categories: disease, drug, pathogen, pathway, protein, gene, small molecule and adverse drug reaction (Figure 2A). For disease nodes, bacterial infections represent the largest group, followed by viral and parasitic infections, reflecting the extensive coverage of clinically infectious diseases (Figure S1A). Pathogen nodes are further classified across major taxonomic groups, including viruses, bacteria, fungi, and parasites, with subcategories (Figure 2B). The quantitative breakdown of these pathogen subcategories (Fig. S1B) reveals insights into their relative representation, such as the predominance of RNA viruses among viral nodes and Ascomycota among fungal nodes.

**Figure 1.**
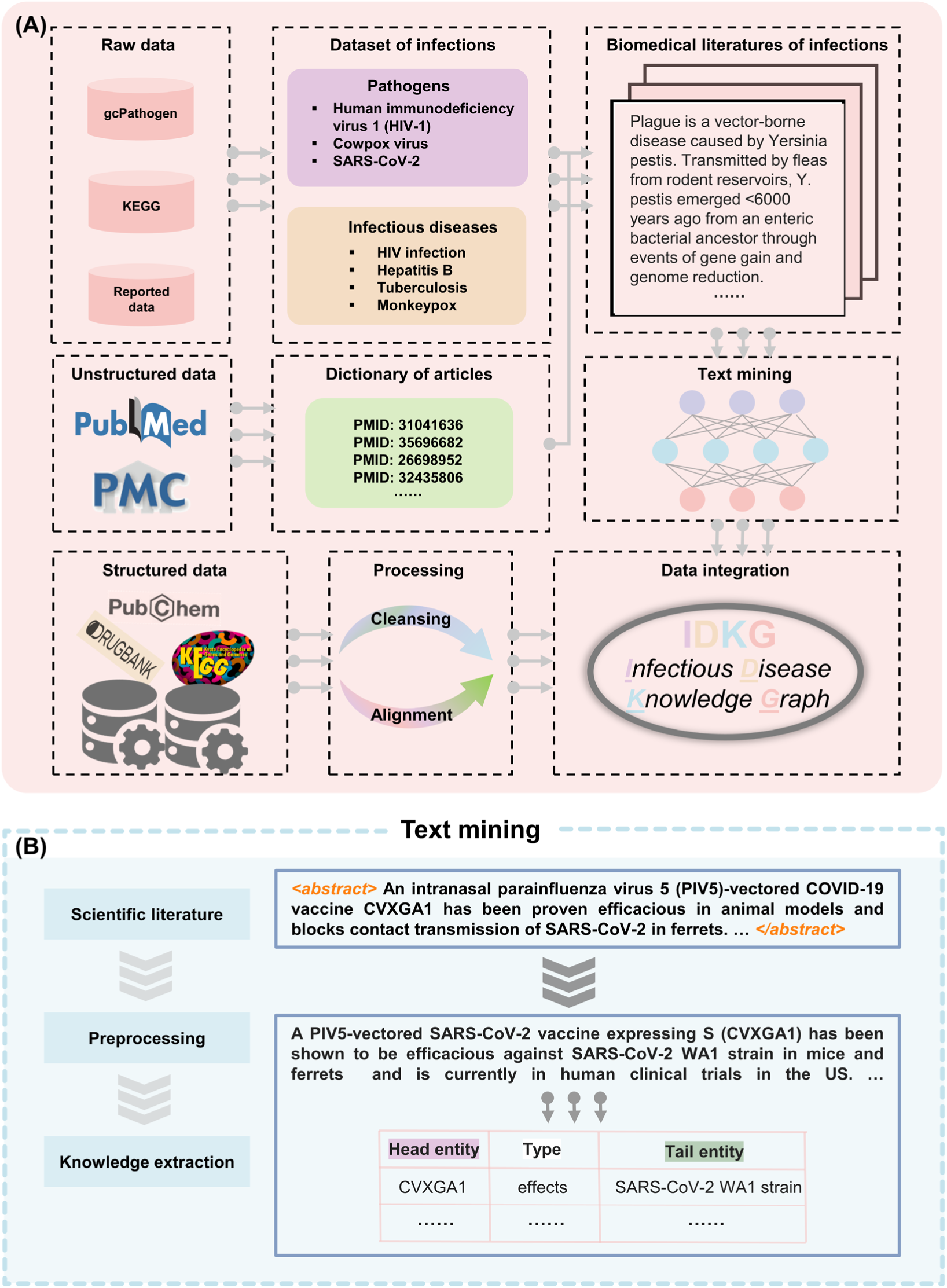
An illustration of IDKG. (A) Pipeline for the construction of the IDKG. First, literature on infections is screened and analyzed using a text mining module; then, biomedical databases from various fields are acquired and processed; finally, these two information sources are integrated to create a comprehensive knowledge graph. (B) The text mining pipeline encompasses the stages of scientific literature acquisition, preprocessing and knowledge extraction.

**Figure 2.**
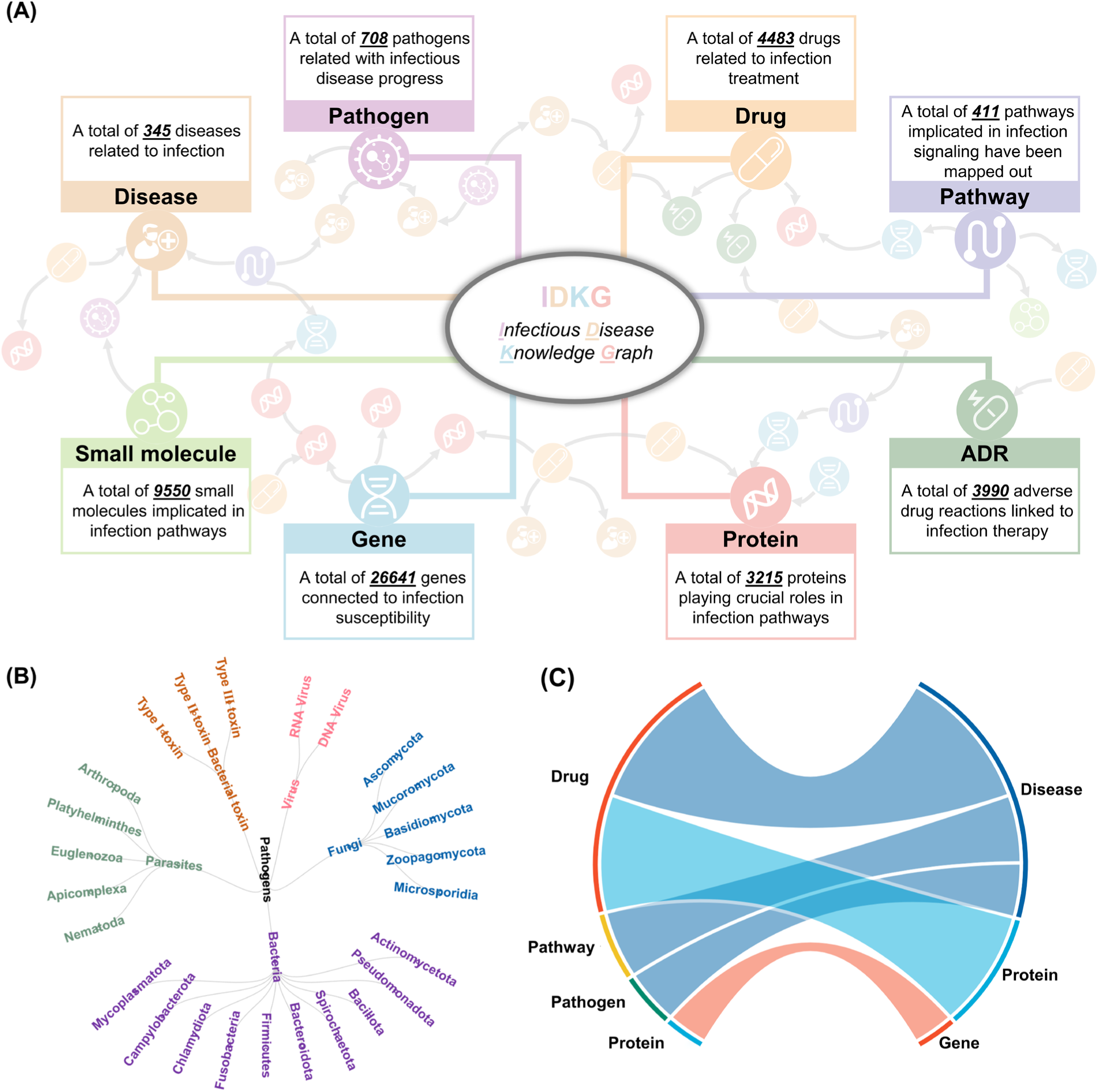
Comprehensive visualization of the IDKG structure and data distribution. (A) Distribution of various node types in IDKG. (B) Taxonomic classification of pathogens in the IDKG, including viruses, bacteria, fungi, and parasites, represented in a hierarchical structure. (C) Chord diagram illustrating the relationships between major node types, such as drugs, diseases, genes, and proteins, emphasizing the interconnected nature of the graph.

Furthermore, the attributes of nodes are designed to be as comprehensive as possible to enhance the scalability of IDKG for downstream tasks (Figure S1C and Table S1). For instance, drug nodes of IDKG include a wide array of chemical informatics and pharmacokinetic properties. These contain the International Union of Pure and Applied Chemistry (IUPAC) name, logarithm of the partition coefficient (logP) values, half-life durations and comprehensive descriptions of the mechanisms through which these drugs exert therapeutic effects.

In addition, we established multiple types of edges to connect different node types, reflecting their biological relationships. An overview of the connectivity patterns among major entities is visualized in Figure 2C, highlighting the intricate interactions between drugs, diseases, proteins, and pathways. Notably, the connections are particularly important, as they form the basis for understanding disease mechanisms, identifying potential drug targets, and evaluating therapeutic strategies.

### Comprehensive medical knowledge interactions via IDKG

In the realm of medical research, the swift and precise retrieval of information is of paramount importance^23^. Researchers can easily acquire specialized cross-disciplinary knowledge through interactions with the IDKG. When investigating HIV infection, researchers can greet with an array of information through keyword queries, which presented as a subgraph (Figure 3A). Then they can further explore the different types of entities within the subgraph to gain more detailed knowledge (Figure 3B). For instance, details on Etravirine—including its IUPAC name, molecular formula, and mechanism—are readily accessible, offering researchers immediate insights into the drug’s therapeutic role. In addition, IDKG integrates crucial data on drug-drug interactions (DDIs), illustrated by Ketoconazole in Figure 3C. This view highlights potential interaction risks, which are further annotated with mechanistic underpinnings (Figure 3D), supporting safer and more effective treatment strategies. More examples can be viewed in Figure S2.

**Figure 3.**
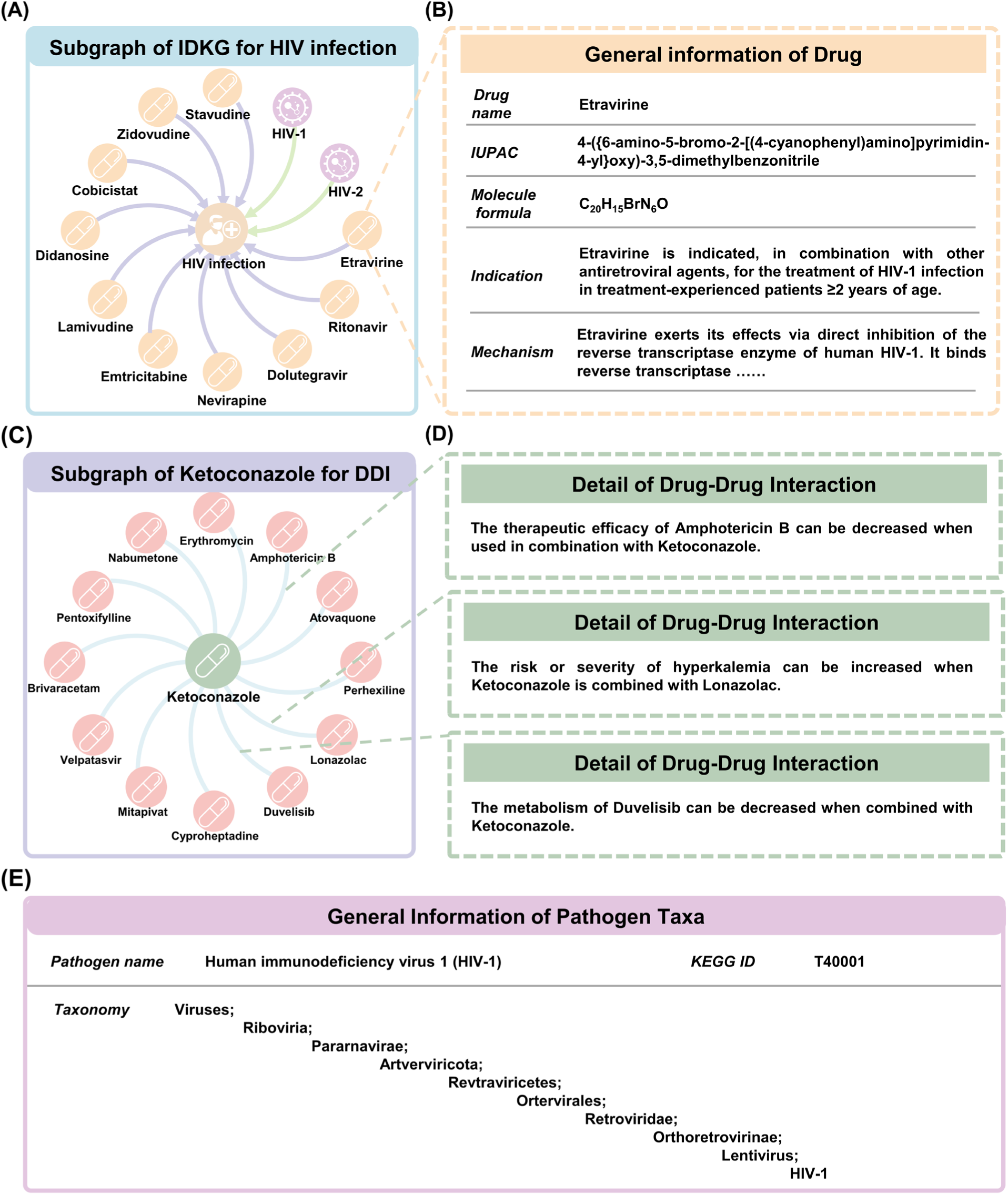
Knowledge interactions with the IDKG. (A) Information associated with HIV infection is presented as a subgraph, including pathogens and drugs. (B) Detailed chemical and pharmacological information for Etravirine, which serves as a representative drug node in the IDKG. (C) Drug-drug interaction (DDI) subgraph exemplified by Ketoconazole. (D) Detailed comments on drug-drug interactions to illustrate potential risks and effects. (E) Basic information about the pathogen (HIV-1), including taxonomic status and important characteristics.

### Node similarity and network topology of IDKG

Similarity analysis in a KG is a useful analytical method that measures the entity-entity or relation-relation similarity^23^. To investigate whether a disease is more likely to be proximal to other diseases that share therapeutic drugs, we compared similarity between disease pairs that did and did not share drugs through graph embedding. Graph embedding methods map each node to a vector by capturing structural information in the graph^24–26^. These embeddings serve as compact and informative representations of the graph’s topology. To determine an optimal embedding dimension that captures all structural information effectively while maintaining parsimony, we employed a normalized embedding loss function^27^. The approach quantified the difference between embeddings across dimensions, ensuring that lower-dimensional embedding remain both effective and computationally efficient.

Our analysis revealed that an embedding dimension of 148 achieves optimal performance with a normalized accuracy threshold of 0.05 (Figure 4A). We then proceeded to investigate the cosine similarity between disease pairs that share common drugs and those that do not under this embedding configuration andvisualized the similarity distribution using kernel density estimation (KDE) to explore how shared therapeutic agents influence the similarity, which was shown in Figure 4B. It revealed distinct patterns between the two groups, providing insights into the relationship between shared drugs and the structural embedding similarity of diseases. For instance, diseases such as HIV infection and hepatitis B, which share antiviral drugs lamivudine, exhibited closer embedding similarities. This reflects their overlapping treatment strategies targeting viral replication pathways. In contrast, diseases such as tuberculosis and malaria, which rely on entirely distinct therapeutic approaches, showed more divergent embeddings.

**Figure 4.**
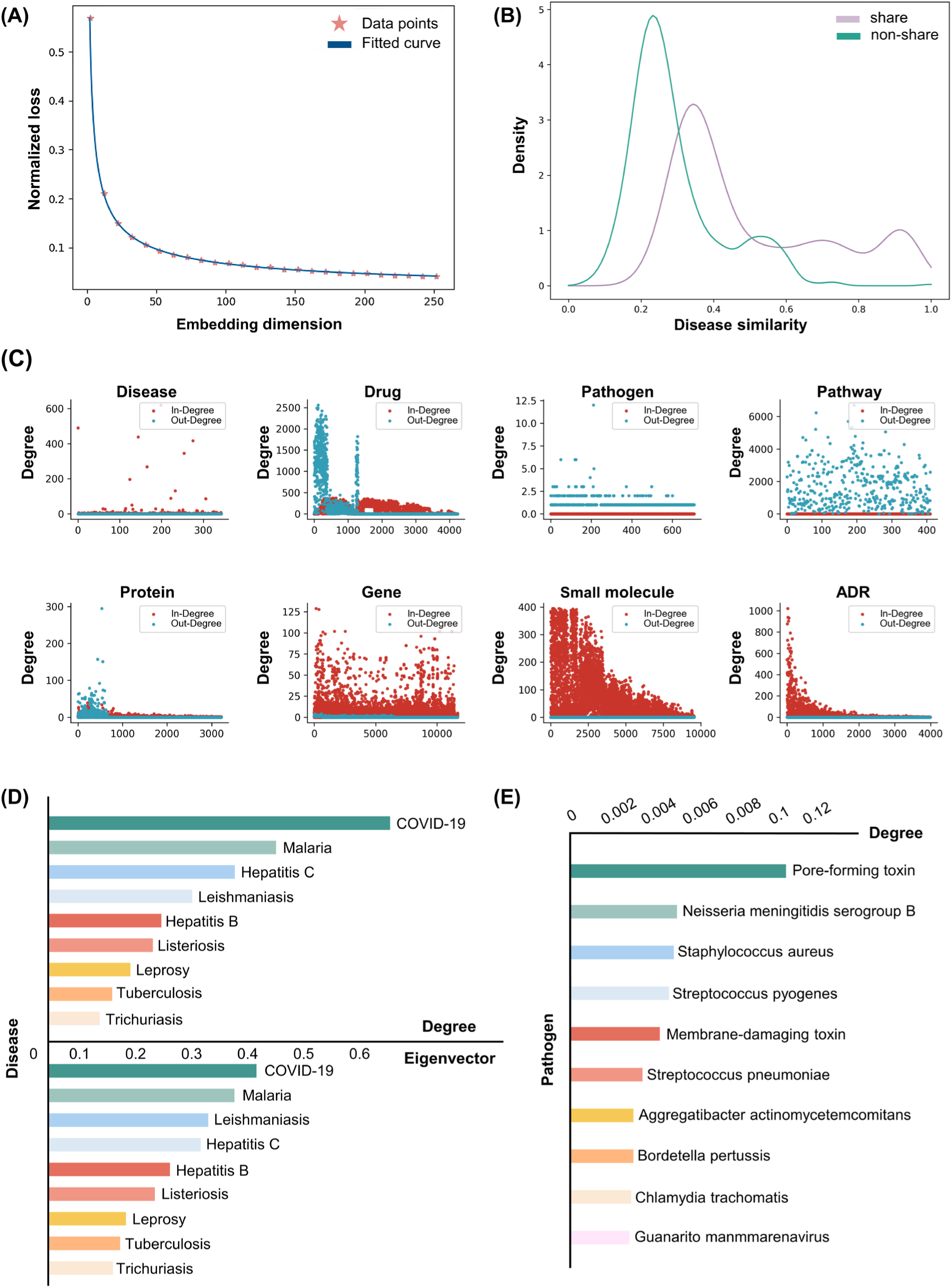
Similarity and topology analysis of IDKG. (A) The relationship between the embedding dimensions and normalized loss. The data points represent the measured normalized loss for each embedding dimension, while the fitted curve highlights the trend. (B) Kernel density estimation (KDE) curves depicting the distribution of cosine similarity between disease pairs. (C) In-degree and Out-degree distribution of nodes in IDKG. (D) Centrality analysis on pathway-disease relationships using degree and eigenvector centrality. (E) Centrality analysis on pathogens using degree centrality.

To further characterize IDKG’s topological properties, we performed degree distribution and centrality analysis. Degree distribution analysis examines the number of connections each node has within the knowledge graph. Analyzing the distribution of these degrees across all nodes provides insights into the overall connectivity patterns and the presence of highly connected hubs within the network. The visualization of in-degree and out-degree patterns is shown in Figure 4C. Such heterogeneity in degree distributions highlights the diverse functional roles and connectivity patterns of distinct entity types within the IDKG, thereby providing a solid foundation for downstream network-based analyses. Centrality analysis provides an effective way to identify the most influential nodes within the network, revealing key contributors to its overall structure and functionality. As shown in Figure 4D, we calculated the centrality metrics for the pathway-disease relationships and analyzed two centrality indicators for the nodes in the graph: degree centrality and eigenvector centrality. The analysis reveals that diseases such as COVID-19, malaria, and hepatitis C exhibit high centrality values, indicating their significant roles in the network. These diseases are highly connected to other nodes, reflecting their importance in the pathway-disease interactions captured in IDKG. By identifying these key nodes, the centrality analysis provides a deeper understanding of the network structure, offering potential directions for exploring disease mechanisms and prioritizing therapeutic targets. Similarly, we also conducted the analysis on the pathogen nodes to highlight their pivotal roles within this field (Figure 4E)^28,29^.

### IDKG for drug repurposing

The development of anti-infective drugs has primarily focused on broad-spectrum antibiotics. However, their overuse may exacerbate antimicrobial resistance. In contrast, effective narrow-spectrum antibiotics can be a good choice for the treatment of infections with drug-resistant bacteria with their targeted activity^9^. Drug repurposing methods based on phenotype screening are time-consuming and inefficient, making it difficult to meet the urgent demand for the rapid development of anti-infective drugs^30^.

To address the limitations, KG-based graph algorithms provide a promising alternative. As shown in Figure 5A, we extracted a subgraph from IDKG which contains all drug, disease and protein entities along with their connecting edges as inputs. To model the potential relationships between these nodes, we have designed two distinct encoder-decoder architectures (Figure 5B). For the encoder, we experimented with two different approaches: (1) Graph Convolution Network (GCN); and (2) a hybrid architecture combining GCN with Graph Attention Network (GAT) to capture features. In the decoder, we utilize Multilayer Perceptron (MLP) layer to predict scores, followed by the application of the Sigmoid function to map these scores into a 0-1 range.To evaluate the performance of our models, we performed 10-fold cross-validation. The area under the receiver operating characteristic curve (AUROC) and the area under the precision-recall curve (AUPRC) have been widely used in bioinformatics research^31,32^, and are adopted to evaluate the overall performance of our models. As shown in Table 1, the final average values of accuracy, AUROC, and AUPRC achieved by M1 are 0.8486, 0.9680, and 0.9749, respectively, while M2 yields values of 0.8470, 0.9629, and 0.9725.

**Figure 5.**
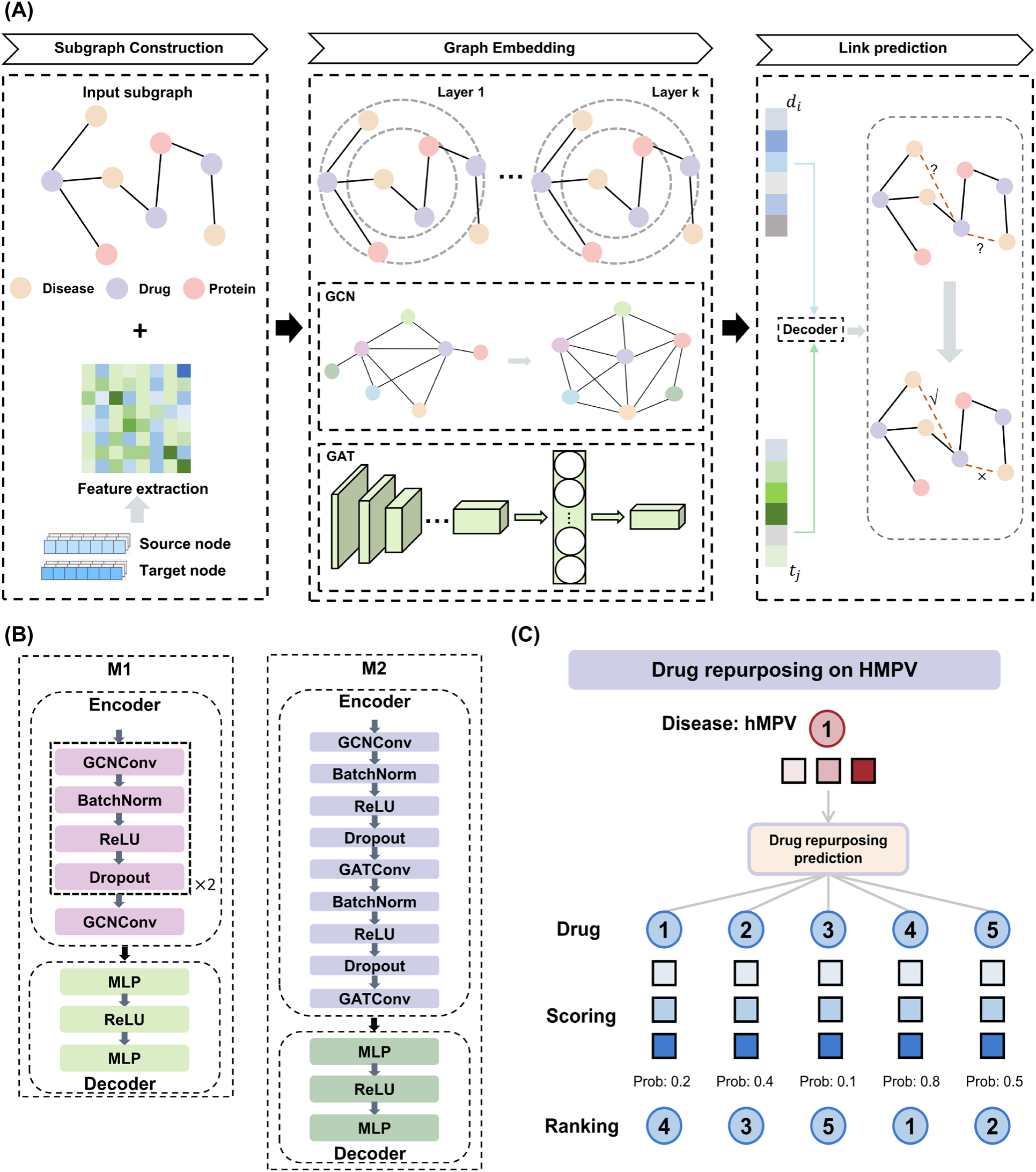
Pipeline for drug repurposing and a case study. (A) The pipeline for prediction of drug repurposing. A subgraph extracted from the IDKG serves as the input for the model, which encodes the intricate relationships between nodes using sophisticated encoders and decoders. This process allows the model to predict potential relationship scores, providing valuable insights to support decision-making processes. (B) Two kinds of encoder-decoder architectures. (C) A case study on hMPV for drug repurposing.

**Table 1.** The values of evaluation metrics obtained under the 10-fold cross-validation.

| Model | Accuracy | AUROC | AUPRC |
| --- | --- | --- | --- |
| M1 | $0.8486 \pm 0.0105$ | $0.9680 \pm 0.0073$ | $0.9749 \pm 0.0038$ |
| M2 | $0.8470 \pm 0.0127$ | $0.9629 \pm 0.0111$ | $0.9725 \pm 0.0088$ |

Human metapneumovirus (hMPV) is considered the most common cause of viral respiratory tract infections in infants. Approximately 90% of children worldwide have experienced at least one hMPV infection by the age of five^33,34^. Currently, there is neither vaccine nor antiviral treatment available on the market^35^. Then, we chose the model with better performance for hMPV drug repositioning study (Figure 5C). The top-15 candidates are shown in Table 2. Among these, ribavirin and emetine have been reported to exhibit anti-hMPV activity^34,36^. Ribavirin, a broad-spectrum antiviral, is known to inhibit RNA synthesis, making it effective against a wide range of RNA viruses, including hMPV^36^. Emetine, traditionally used as an anti-protozoal agent, has shown potential antiviral activity in cell-based assays. For example, a study demonstrated that Emetine inhibits hMPV-mediated GFP expression in RPE cells, suggesting its potential as an antiviral agent against hMPV^34^. These findings align with their high rankings in our prediction results, further validating the reliability of our model.

**Table 2.**
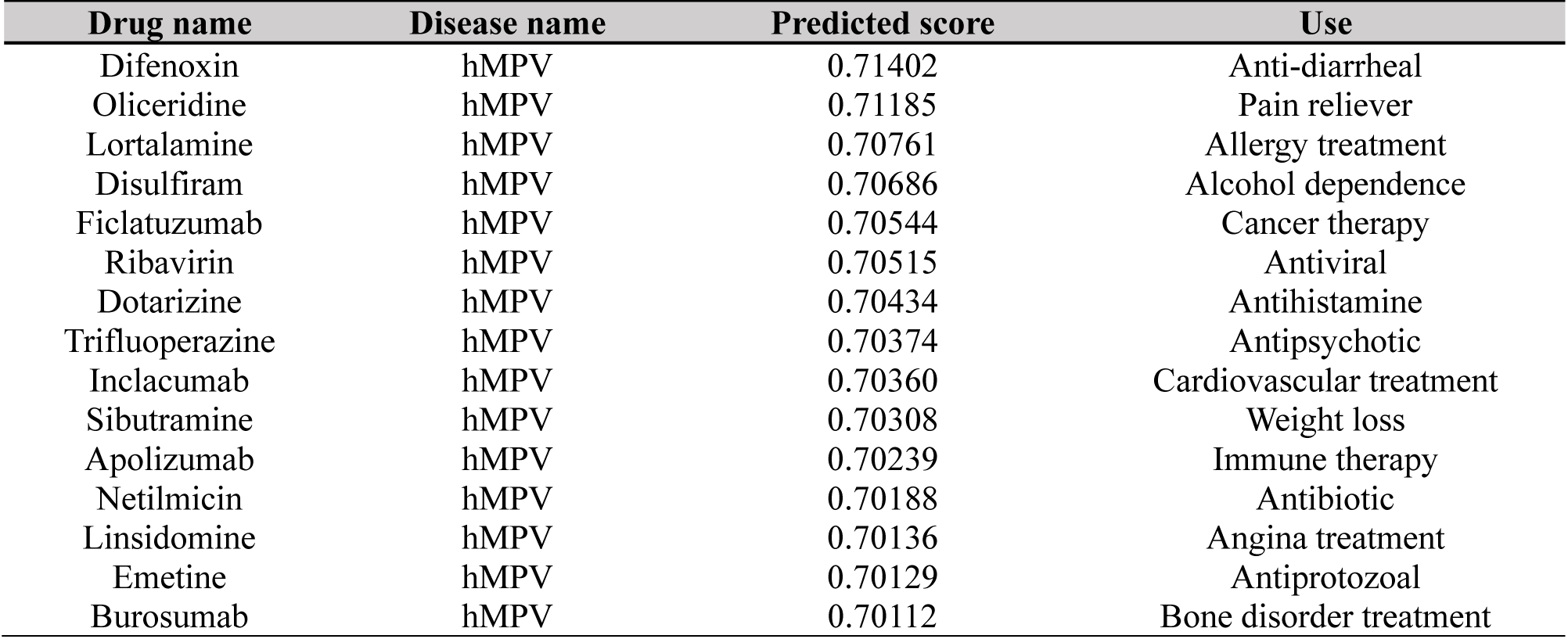
Summary of the top 15 predicted drug-hMPV pairs.

To further expand the downstream applications of the IDKG, we performed link prediction across all drug-disease pairs. Then, we selected M1 for inference to identify the top 10 highest scoring drug-disease pairs (Table S2). The predicted drugs are mostly not traditional anti-infective agents; instead, they may exert their effects through immune modulation or indirect mechanisms to inhibit specific pathogens. For example, vismodegib, used in cancer treatment, may indirectly influence immune responses, potentially affecting the body’s ability to fight infections in immunocompromised patients. Lefamulin, primarily used for pneumonia^37^, works by inhibiting bacterial protein synthesis, but its effects on bacterial virulence and the host immune response may also contribute to infection control. This non-conventional therapeutic approach offers a novel perspective, particularly in addressing infections caused by multidrug-resistant pathogens. Further research into the mechanisms of action and potential expansion of clinical indications for these drugs could pave the way for innovative strategies in the treatment of challenging infectious diseases.

## Discussion

Humanity continues to face the severe threat of infectious diseases. Due to challenges such as the high variability and drug resistance of pathogens, researches on many infectious diseases remain in its early stages or even lack a comprehensive understanding. Nevertheless, the volume of infectious disease related literature has grown substantially, encompassing a wealth of fragmented information scattered across various sources (Figure S3). This vast and rapidly expanding corpus holds untapped potential for uncovering novel insights and advancing our understanding of infectious diseases. However, effectively extracting and synthesizing this knowledge remains a formidable challenge, highlighting the urgent need for innovative approaches to systematically organize and leverage this information.

In response to these challenges, we developed the IDKG. Through the integration and standardization of multimodal data, the IDKG comprises nearly 50,000 nodes (eight types) and an extensive network of over 1.2 million edges (11 types). It consolidates information on diseases, pathogens, drugs, and related biomedical entities, providing a unified framework for advanced research. Compared to existing biomedical knowledge graphs, which are often focused on specific domains or limited in scope^23,38–42^, our IDKG is unique in its breadth and adaptability to the infectious disease landscape.

Building on its high-quality data structure, the IDKG serves as a powerful knowledge base that can be seamlessly integrated with emerging AI technologies. This fusion enables more efficient and accurate exploration of infectious disease mechanisms, facilitating the identification of novel drug targets, new therapeutic mechanisms, and even repurposing existing drugs for new indications. By leveraging advanced machine learning models, such as deep learning and network-based algorithms, the IDKG can uncover hidden patterns and predict potential drug-disease interactions, thereby accelerating the development of anti-infective therapies. This integration not only enhances the speed of drug discovery but also offers insights into innovative strategies for combating infectious diseases and addressing antimicrobial resistance.

However, in the context of drug repositioning research, which involves predicting new relationships between drugs and diseases, the link prediction task is central to completing and refining the knowledge graph. From an informatics perspective, enhancing the prediction of potential connections within the graph remains a key challenge. The combination of LLMs and Retrieval-Augmented Generation (RAG) offers a significant advancement. The powerful semantic representation capabilities of LLMs, complemented by the retrieval function of RAG, enable the discovery of richer and more accurate connections. This synergy not only enhances the model’s ability to understand and reason within multidisciplinary knowledge domains but also facilitates the prediction of previously unknown links within the knowledge graph^43,44^. As a result, this approach holds the potential to improve the completion and structure of the graph, driving more effective and insightful drug repositioning research. IDKG constructed in this study can be integrated with LLMs to further enhance the model’s reasoning capabilities. By combining IDKG with LLMs, we can better uncover and predict new, relevant knowledge related to infectious diseases, thereby advancing our understanding and discovery in this critical field.

## Methods

### Data architecture

The data architecture of IDKG is meticulously designed to incorporate both structured and unstructured data, ensuring a comprehensive representation of biomedical knowledge. The structured data component is built by integrating high-quality resources from diverse domains. In parallel, the unstructured data component focuses on extracting knowledge from infection-related literature, curated from the vast repositories of PubMed (https://pubmed.ncbi.nlm.nih.gov) and PubMed Central (https://www.ncbi.nlm.nih.gov/pmc).

### Structured data processing

We utilized a series of authoritative biological databases to construct a comprehensive dataset as part of the development of the IDKG. The dataset provides extensive coverage of biomedical entities such as disease, drug, gene and pathogen. Using the API provided by the World Health Organization (https://icd.who.int/icdapi), we retrieved the ICD-11^45^ to compile a dictionary of all recognized infectious diseases, thereby delineating the scope of the study. Subsequently, biomedical entities and their associated biological relationships were extracted and filtered from databases including UniProt^46^, HGNC^47^, DrugBank^48^, CTD^49^, KEGG^50^, and PubChem^51^, referencing the disease dictionary as a reference.

To ensure data consistency and integrity, the raw data underwent a series of cleaning and preprocessing steps. First, duplicate entries were identified and systematically removed based on unique identifiers. Missing values (NaN) were addressed by imputing data from cross-referenced sources or by removing incomplete entries when imputation was not feasible. Entity names and relationship terms were standardized using controlled vocabularies to harmonize nomenclature across databases. The result was organized into CSV files for further processing.

### Text mining from biomedical literature

Text mining aims to obtain hidden relationships in unstructured text and permit the generation of new knowledge^52^. It primarily consists of two key components: data selection and processing, and knowledge extraction. The pipeline is shown in Figure 1B.

### Data selection and processing

The original data is mainly from PubMed abstracts and PubMed Central full-text articles. To date, more than 34 million PubMed and nearly 2.8 million PMC articles can be available for free download in XML format. In short, XML documents use tags to describe the structure and elements within the data. The article’s basic information is stored in the tag pairs, such as title, abstract, keywords, DOI, and so on. We extracted fundamental data elements such as the PMID (PMCID), title, abstract, and keywords from each article, which serve as the basis for identifying infection-related literature, by utilizing the Python ElementTree module in conjunction with XPath.

In addition, we constructed a comprehensive dictionary of pathogens and infectious diseases by integrating resources from KEGG^50^, gcPathogen^53^, and a table of bacterial pathogens reported as of 2021^54^. Subsequently, string-matching algorithms were employed to identify all article IDs related to infectious diseases. The abstracts and full texts of these articles were then retrieved and processed to establish a specialized text corpus database for knowledge extraction, comprising over 2.3 million pieces of literature.

### Knowledge extraction

In the field of Natural Language Processing (NLP), the multi-head attention mechanism featured in the Transformer^55^ architecture has proven to be a powerful tool for extracting information from text. Pre-training models have demonstrated their powerful capability in NLP. On the GLUE benchmark, a widely used benchmark for natural language understanding, pre-training-based methods outperform non-pre-training methods by a large margin^56^. With the success of pre-training in general NLP, people explore adapting pre-training models into the biomedical domain. There are two main kinds of foundation models: BERT^57^ and GPT^58–60^, mainly for language understanding tasks and language generation tasks respectively.

Although fine-tuned models such as BioBERT^61^, PubMedBERT^62^, BioGPT^63^ have demonstrated impressive performance in tasks such as Named Entity Recognition (NER) and Relation Extraction (RE) within the biomedical domain, they are primarily specialized in a single aspect and do not effectively balance comprehension with generation capabilities^63^. To address the limitations, we utilized REBEL^64^, a seq2seq model based on BART^65^ that performs end-to-end relation extraction to extract all hidden relationships from the prepared corpus of literature related to infection. The extracted triples were then annotated and refined by filtering out data where both the head and tail entities were biomedical entities, yielding the final dataset.

### Graph database and data integration

Graph databases are NoSQL databases that represent and store data using graph structures. The graph structure is a collection of nodes and edges that represent relationship between the nodes and properties. Storage of data in such a structure facilitates access to densely connected data by providing graph traversal linear times^66^. The database is built using Neo4j (https://neo4j.com), which provides a query language specific for graph structures, Cypher, and an extensive library of procedures and functions (such as APOC library and the Graph Data Science library) that can be used for data integration and analysis.

We selected diverse node labels and relationship types to design the graph data model. These nodes and relationships were defined based on the types of biological entities and their interactions. For each node label, identifiers were assigned using widely recognized biomedical ontologies or standard terminologies. Ontologies represent concepts, such as nodes (e.g., disease), and provide an acyclic graph structure that describes their relationships. However, some nodes in the graph were not covered by existing ontologies and required standardization using identifiers from selected biomedical databases (e.g., UniProt for proteins, DrugBank for drugs). These identifiers were then mapped to the corresponding nodes in the graph, ensuring consistency and interoperability across various data sources. Once the graph data model was established, we imported the curated data into Neo4j, utilizing Cypher queries to create nodes and relationships according to the predefined schema. Then we got the knowledge graph database.

## Data availability

The authors declare that all data supporting the findings of this study are available within the article or from the corresponding author upon reasonable request.

## Code availability

The code package is freely available via GitHub at https://github.com/henryFan128/IDKG.

## Acknowledgements

This work was supported in part by the National Natural Science Foundation of China (82425104 and 82150208 to H.L., 82173690 to S.L.), the National Key R&D Program of China (2022YFC3400501, 2022YFC3400504), S.L. is also sponsored by the Shanghai Rising-Star Program (23QA1402800).

## Author contributions

Conceived the study: H.L. and S.L. Collected the data: H.F., F.L. and Y.D. Processed the data and implemented the architecture: H.F., L.G. and Y.X. Wrote the paper: H.F. Revised the paper: H.F., Y.Z. and S.L. All authors have read the final manuscript and approved it for publication.

